# Fatty Acid Synthase Binds Cellular mRNAs and its Depletion Reduces Translational Efficiency

**DOI:** 10.64898/2026.11.05.764384

**Authors:** Yelena Perevalova-Pinzul, James Marks, Marcin Sajek, Stephen Clarke, Samantha Lisy, Neelanjan Mukherjee, Sezen Meydan, Manuel Ascano

## Abstract

Metabolic enzymes with non-canonical RNA binding activity have drawn a growing interest, due to their implications for basic biology and disease. Here we present the first RNA interactome of fatty acid synthase (FASN), an emerging oncological and virological target. Utilizing crosslinking-based approaches, we characterize the direct RNA association of FASN in two human cell lines, identifying over 6000 binding sites across over 3500 protein-coding genes, with a striking preference for the 5’ UTR. We provide a functional cross-screen on the effects of FASN depletion on cellular translation. This integration with ribosome profiling indicated a preferential reduction in the translational efficiencies of FASN-associated mRNAs – an effect shown to be dependent on binding location and site number. Associated transcripts were enriched for morphogenic and cell division-related functions, as well as palmitoylated proteins. Together, these findings identify a transcript-selective, position-dependent relationship between FASN RNA binding and translation, expanding the cellular functions attributable to this metabolic enzyme.

## INTRODUCTION

RNA binding proteins (RBPs) function as central regulators of co- and post-transcriptional gene expression, controlling processes including RNA processing, localization, stability, and translation. Large-scale studies of cellular RNA-protein interactions have substantially expanded the recognized RBP repertoire, revealing numerous proteins without canonical RNA-binding domains or previously established functions in RNA biology, including many metabolic enzymes^1–6^. For many of these non-canonical RBPs, however, the biological significance of RNA binding remains elusive, particularly when RNA interaction must be distinguished from the protein’s established enzymatic or metabolic function^7,8^.

One such protein is fatty acid synthase (FASN). FASN is a metabolic enzyme best characterized for its role in *de novo* fatty acid synthesis. FASN functions as a homodimer^9^ and is the major cellular source of longchain fatty acids, principally palmitate^10^. FASN has been implicated in diverse physiological and pathological processes, including development and nervous system function, cognition, cell division, oncogenesis, and immune responses^11–19^. Much of this biology has been interpreted through the canonical lipogenic activity of FASN, and consequently considerable effort has focused on pharmacological inhibition of its catalytic activity^10,20–23^. However, accumulating observations suggest that some functions of FASN may not be explained solely by its catalytic activities, raising the possibility of additional roles.

Evidence for such non-canonical functions has emerged particularly from studies of viral infection and innate immunity. FASN has been identified as a host interactor for multiple viruses^24–26^ and has been found in direct association with viral RNAs, including influenza^27^ and SARS-CoV-2^28^ genomes. Although the lipogenic activity of FASN contributes to the replication of several viruses^29,30^, other observations suggest functions that are at least partly separable from its catalytic activity, including effects on viral polymerase processivity^26,31^. More recently, FASN was reported to bind endogenous double-stranded RNAs and modulate cellular responses to exogenous dsRNA^17^ – further supporting RNA interaction as a biologically relevant property of FASN.

During chikungunya virus infection, FASN can promote viral translation even in the presence of a potent FASN catalytic inhibitor^27^. A relationship between FASN and translational regulation has also emerged outside infection: in diffuse large B-cell lymphoma FASN promotes an S6K1-dependent translation program that supports oncogenic proliferation^32^. Together, the convergence of RNA binding and translational phenotypes across distinct biological contexts suggests that interaction with RNA represents a broader cellular function of FASN linked to post-transcriptional regulation.

Consistent with these observations, FASN has repeatedly been identified in large-scale studies of the cellular RNA binding proteome^2,3,33–39^, yet its cellular RNA interactome has not been systematically characterized. Although FASN has been implicated in translational regulation in several biological contexts^27,32^, whether its cellular RNA interactions are associated with transcript-specific effects on translation remains unknown. We therefore sought to define the endogenous RNA interactome of FASN and determine whether FASN-associated mRNAs exhibit altered translational regulation following FASN depletion.

## RESULTS

### Fatty Acid Synthase is a Bona Fide RNA Binding Protein

Given prior work and observations, we hypothesized that FASN is capable of a direct interaction with RNA in a biologically relevant manner. To test this possibility, we performed crosslinking and solid-phase purification (CLASP) in order to determine whether FASN could be crosslinked to RNA (Figure 1A). Metabolic labeling of RNA with 4SU allows us to perform stringent RNA pulldowns, after crosslinking at UV wave-lengths of 365 nm, under protein denaturing conditions, limiting capture to direct RNA-protein interactions within a 3 Å radius^40^. After isolation of the crosslinked ribonucleoproteins, we discovered that FASN was enriched in both HEK293 (Figure 1B) and U2-OS cells^27^. Given this RNA-specific binding in the presence of 4SU (Figure 1C), we aimed to identify the cellular RNA interactants of FASN.

**Figure 1.**
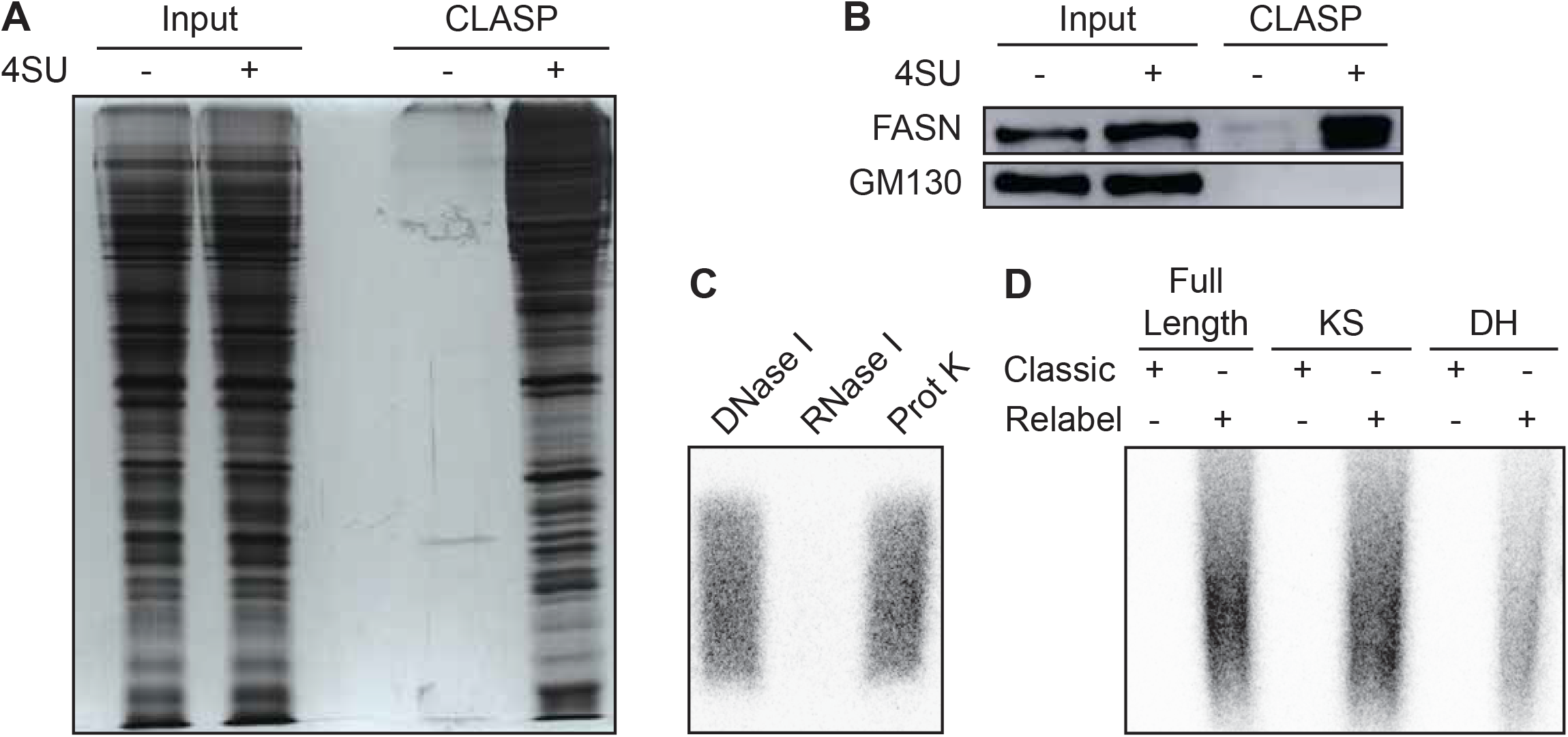
Fatty Acid Synthase contains multiple RNA binding domains which bind to a range of RNA subpopulations. (A) Isolation of the HEK RBPome via CLASP. (B) Detection of FASN enrichment by Western blot. (C) Enzymatic digestion of radioactive products isolated in PAR-CLIP prior to library construction. (D) RNA immunoprecipitated with overexpressed ketoacyl synthase (KS) or dehydratase (DH) domain fragments of FASN.

### PAR-CLIP of FASN

We therefore used photoactivatable ribonucleoside-enhanced crosslinking and immunoprecipitation (PAR-CLIP) to identify cellular RNAs associated with FASN. Because conventional PAR-CLIP conditions yielded insufficient recovery of FASN-associated RNA, we adapted the procedure to improve RNA recovery by incorporating additional radiolabeling procedures (Figures 1D, S1). These modifications enabled recovery of RNA suitable for transcriptome-wide mapping of FASN-associated binding sites. Together with the CLASP experiments, these data establish that FASN associates with cellular RNA. We further found that the ketoacyl synthase (KS) and dehydratase (DH) domains can independently interact with RNA (Figure 1D) consistent with previous observations implicating these domains in RNA interactions^3,27,33,41^.

### Fatty Acid Synthase Reproducibly Binds a Set of Cellular RNAs

Given the capacity of multiple FASN domains to interact with RNA, we next used PAR-CLIP in U2-OS and HEK293 cells expressing FLAG epitope-tagged full-length FASN (FLAG-HA FASN) to identify the RNA binding landscape of FASN. RNA recovered from a band isolated at above 270 kDa, the expected molecular mass of FASN monomers, was subjected to sequencing. At least 100 million reads were obtained from each cDNA library, with approximately 5.2 to 17.3 million uniquely mapped reads per library (Table S2). Across datasets, binding clusters identified by positive T-to-C crosslinked signatures showed high conversion specificity, supporting efficient recovery of FASN-associated RNA; the average T-to-C fraction score was 90% yielding approximately 10-fold enrichment of conversion events at identified clusters.

We identified 15,715 and 11,546 RNA binding sites with at least two unique T-to-C positions in U2-OS and HEK293 cells, respectively. Approximately half of these sites (8,105 and 6,063 in U2-OS and HEK293, respectively) localized to pre-mRNA annotated regions, corresponding to 4,614 and 3,779 genes in U2-OS and HEK293, respectively. Most identified binding sites mapped to RNA polymerase II-derived transcripts (Figure 2A), with approximately half assigned to mRNA, indicating that enrichment of these transcripts in the FASN RNA interactome falls beyond the scope of what would be expected from mRNA abundance alone. This distribution was similar between cell types and remained stable with increasing T-to-C conversion stringency (Figure 2A, 2B, S2). More than half of all ENSEMBL-annotated transcripts identified in one cell type were also detected in the other (Figure S2), demonstrating that FASN associates reproducibly with a broad population of cellular RNAs across cell contexts. Interestingly, over half of annotated transcripts had a singular binding site attributed to them (Figure S2).

**Figure 2.**
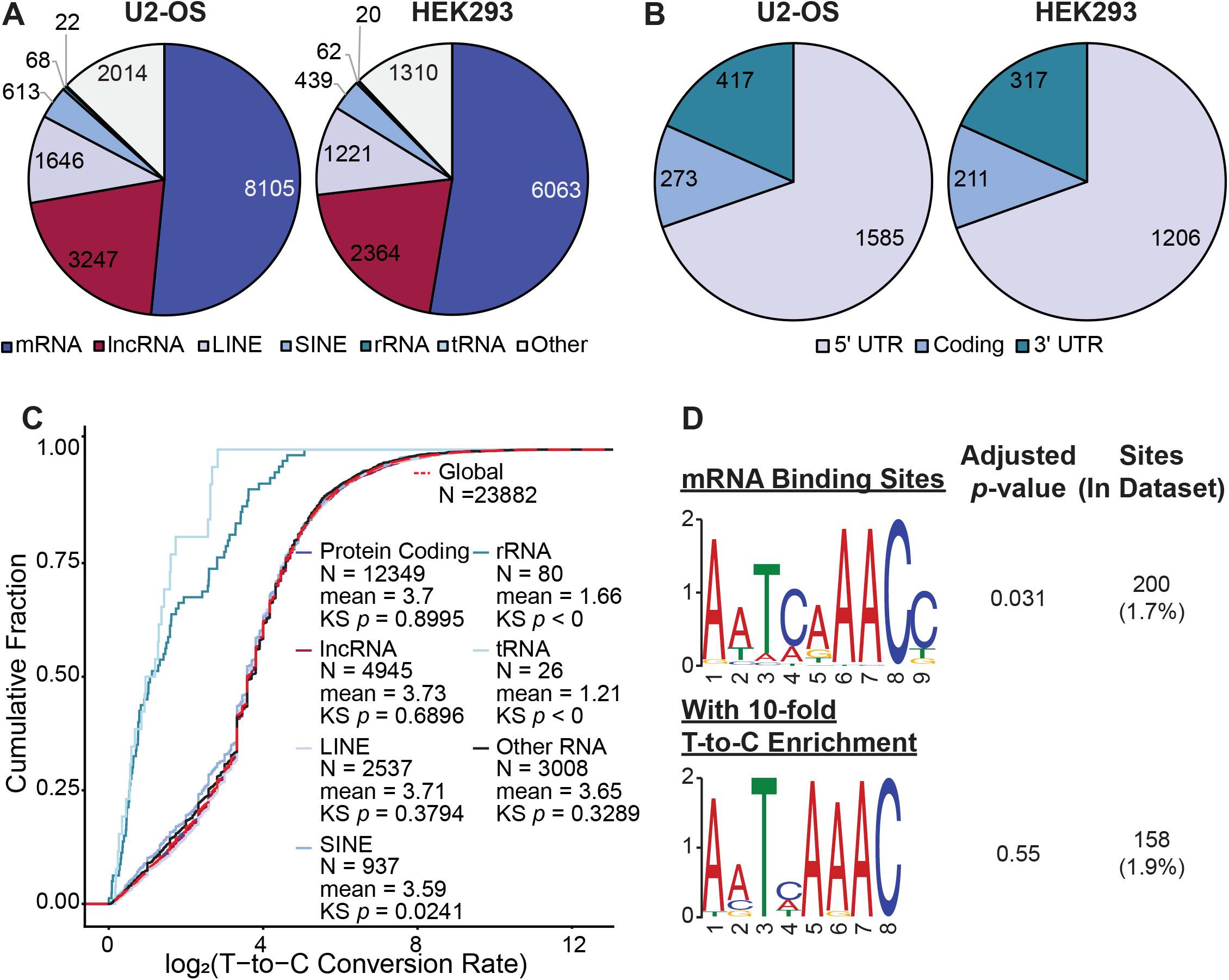
Fatty Acid Synthase interacts with a wide range of Pol-II mediated transcripts. (A) Binding sites identified via PAR-CLIP with a positive T-to-C enrichment value (*n* = 2 biological replicates per line). Statistical similarity: χ2(6, N = 27261), *p* ≈ 0.26. (B) Cumulative distribution analysis of T-to-C transitions in PAR-CLIP targets mapping to annotated RNA. (C) Localization of mature mRNA binding sites identified via PAR-CLIP with a positive T-to-C enrichment value (*n* = 2 biological replicates per line). Statistical similarity χ2(2, N = 4009), *p* ≈ 1.00. (D) Top RNA binding motif identified in unionized PAR-CLIP data.

### mRNA FASN Binding is Enriched in 5’ UTRs

We next asked whether FASN binding was distributed uniformly across mature mRNAs or showed positional enrichment. Among FASN-associated sites mapping to mature mRNAs, more than 60% localized to the 5’ UTR (Figure 2C, S2). This distribution differs markedly from those reported across broader CLIP datasets, in which the 5’ UTRs typically account for a relatively small proportion of identified mRNA binding sites^34^. Thus, FASN exhibits a pronounced preference for the 5’ UTR of mRNAs. Next, we asked whether this positional preference could be explained by a common short sequence motif. Although one statistically significant motif was detected, it was present in fewer than 2% of binding sites and did not remain significant with increasing T-to-C conversion stringency (Figure 2D).

### Fatty Acid Synthase Depletion Reduces Translational Efficiency of Bound Transcripts

Having established that FASN associates extensively with cellular mRNAs and is enriched at 5’ UTRs, we next asked whether FASN-bound transcripts were preferentially affected at the level of translation following FASN depletion. FASN has previously been implicated in translational regulation in both oncogenic and viral contexts^27,32^, providing a basis for testing whether this activity extends to its cellular mRNA targets. We therefore used ribosome profiling together with matched RNA sequencing to determine the changes in translational efficiency following siRNA-mediated depletion of FASN. Due to *FASN*’s status as an essential gene^12,42^, we used RNAi to knock down its expression an experimental approach. Significant reductions in FASN mRNA and protein were confirmed following siRNA treatment (Figure 3A), and translational efficiency was calculated from ribosome-protected fragments normalized to matched RNA-seq measurements (Figure 3 B-E, S3A).

**Figure 3.**
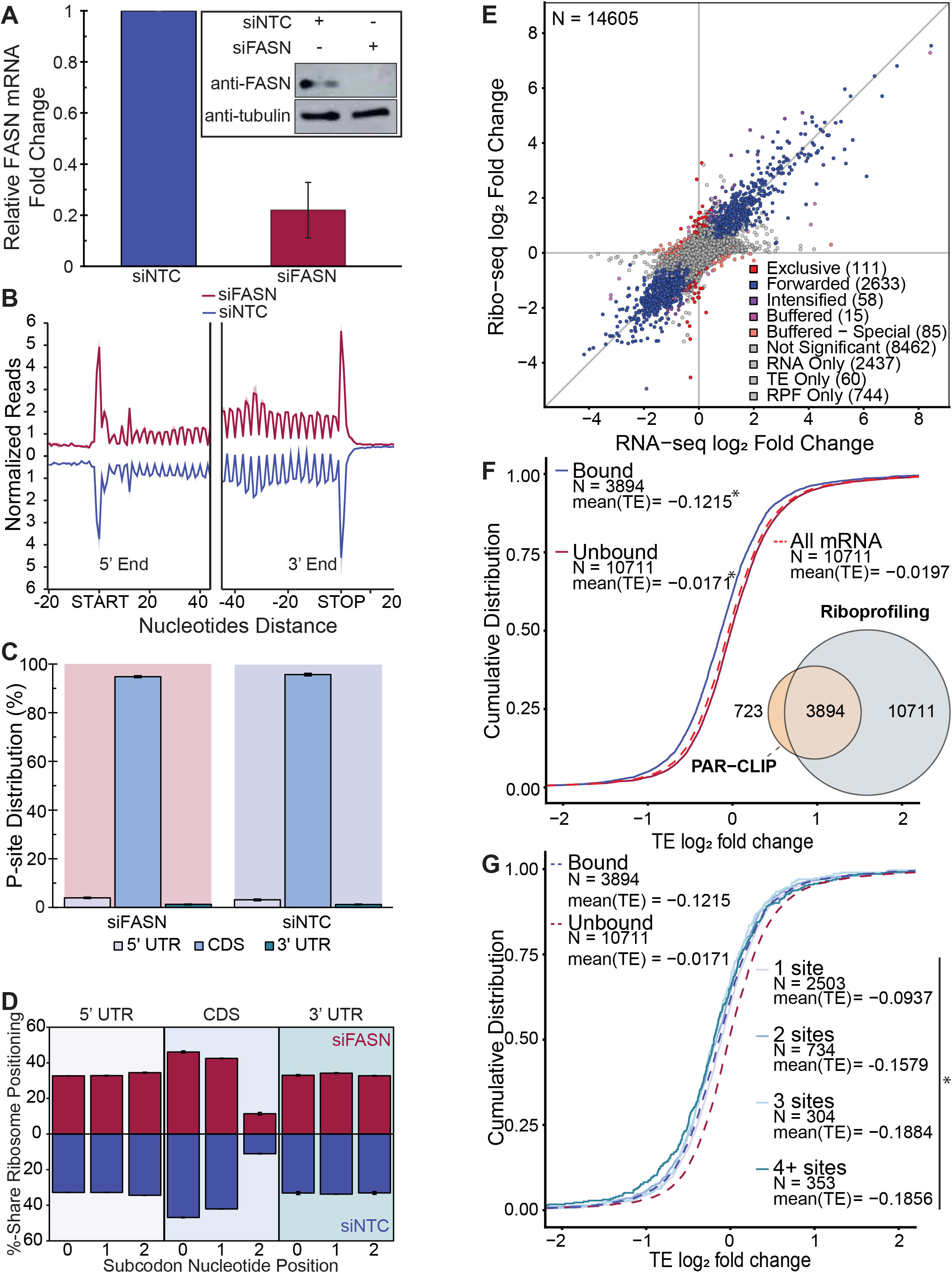
Effects of Fatty Acid Synthase on translation. (A) Knockdown of FASN by siRNA characterized by both relative mRNA levels (*n* = 3 biological replicates) and protein detection via Western blot (inset). (B) Metagene analysis for the distribution of average ribosome density aligned at the start and stop codons (*n* = 2 per condition). (C) P-site distribution of ribosome protected fragments (RPF) in cells undergoing FASN knockdown in U2-OS (*n* = 2 per condition). (D) Subcodon positioning of the P-site (*n* = 2 per condition). EA) Protein-coding transcript RNA expression and RPF enrichment in cells undergoing FASN knockdown in U2-OS (*n* = 2 per condition). (F) Cumulative distribution analysis of translational efficiency changes for all protein-coding transcripts with measurable RPF and RNA-seq signal, separated by the presence of a FASN binding site. Inset: Overlap of annotated transcripts identified as statistically significant riboprofiling hits and genes containing a PAR-CLIP-identified binding site. (G) Cumulative distribution analysis of translational efficiencies of mRNA targets by number of FASN binding sites. * - statistical significance (refer to Table S1)

We next asked whether FASN-bound transcripts were preferentially represented among transcripts with altered translational efficiency following FASN depletion. Across all protein-coding transcripts with measurable ribosome protected fragment (RPF) and RNA-seq signal, FASN-associated transcripts showed a significant reduction in translational efficiency relative to transcripts lacking detectable FASN binding (un-bound) (Figure 3F). This shift was evident across the full distribution, rather than being driven by a small number of strongly affected genes, and was similarly observed when the analysis was restricted to transcripts with individually significant changes (as defined by Chothani and colleagues^43^, Figure S3 B-E). Within that set, transcripts showing changes in translational efficiency independent of RNA abundance were identified alongside those with proportional change in RNA and ribosome-protected fragments. Transcripts whose binding sites were restricted to the coding sequence instead were statistically distinct from all other bound transcripts, showing higher translational efficiencies (Figure S3F), while transcripts bound within non-protein-coding regions were statistically indistinguishable from the general bound population. Furthermore, an increase in the total number of binding sites was associated with a statistically significant decrease in translation efficiency, with transcripts containing more FASN binding sites being less efficiently translated (Figure 3G). Additionally, while T-to-C conversion metrics did not yield a significant decrease, a tendency towards lesser means of translational efficiency was noted (Figure S3G). Overall, FASN-associated transcripts exhibit a preferential reduction in translational efficiency following FASN depletion compared with transcripts lacking detectable FASN binding, with greater reductions observed in those with a multiplicity of binding sites. By contrast, FASN overexpression did not produce an appreciable global change in translational efficiency (Figure S3H).

Among mRNA transcripts exhibiting reduced translational efficiency following FASN depletion, encoded proteins associated with cell division and mitotic structures were significantly overrepresented (Figures 4A and B), consistent with previous observations linking FASN depletion to downregulation of cellular proliferation and cell-cycle progression^15,16,44^. When analysis was restricted to protein-coding transcripts containing at least one FASN binding site, enrichment extended to functions associated with cell division, morphogenesis (including axonogenesis), and cellular mobility, as well as localization to membrane-associated structures (Figures 4C and D). These enrichments were also consistent with the functional categories represented among FASN PAR-CLIP targets when examined independent of ribosome profiling analyses (Figure S2) and consistent with FASN’s documented role in development of neurological tissues^11–13^ and as a mediator of membrane targeting^45,46^.

**Figure 4.**
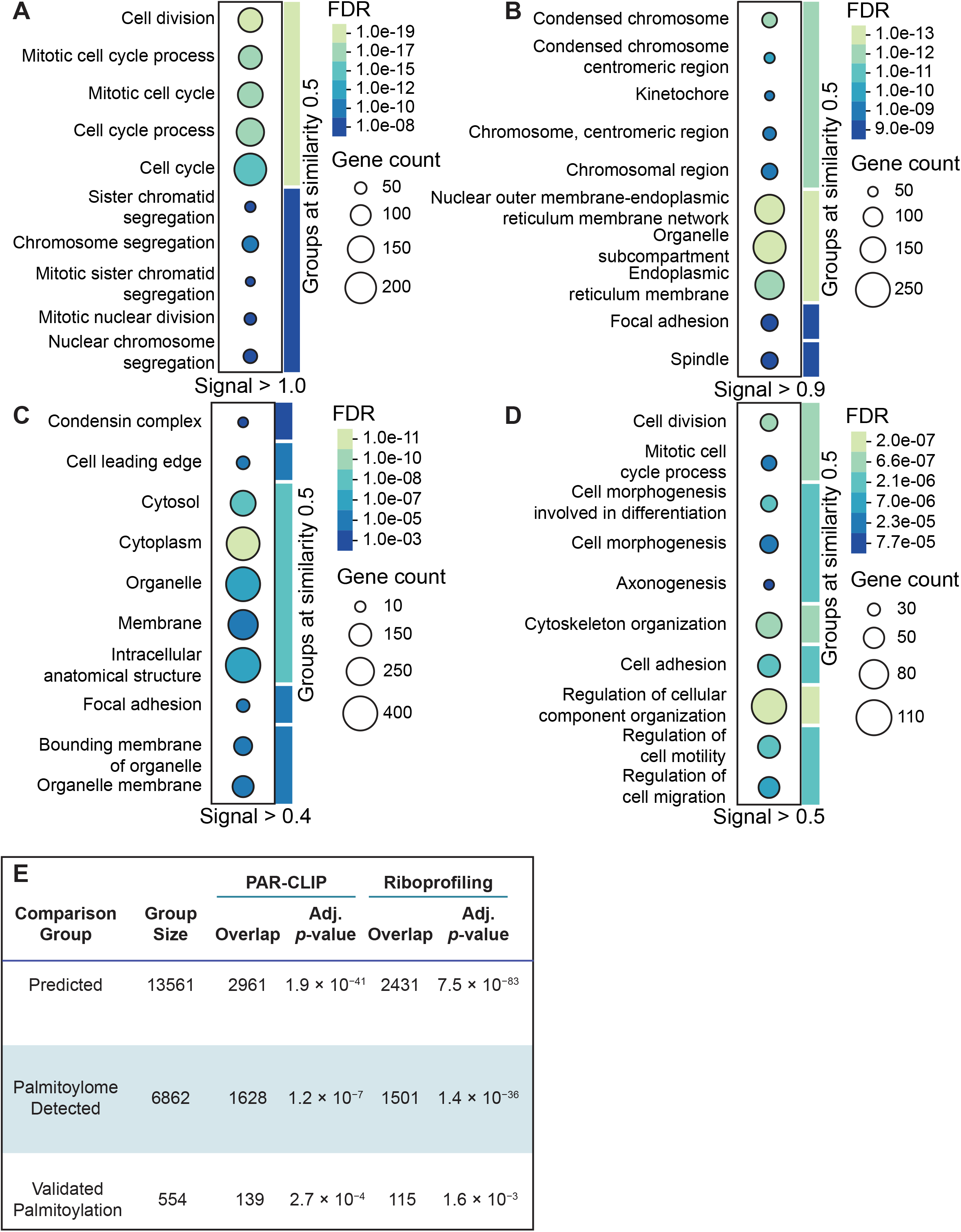
Ontological analysis and palmitoylation status of FASN-regulated genes. (A) Gene ontology analysis of protein-coding hits with negative TEs, querying biological processes. (B) Gene ontology analysis of protein-coding hits with negative TEs, querying localization to cellular compartments. (C) Gene ontology analysis of protein-coding hits with negative TEs and containing at least one FASN binding site, querying biological processes. (D) Gene ontology analysis of protein-coding hits with negative TEs and containing at least one FASN binding site, querying localization to cellular compartments. (E) Overrepresentation of palmitoylated proteins in U2-OS PAR-CLIP and riboprofiling (FASN-bound) datasets. Predicted – computationally predicted to be palmitoylated. Palmitoylome detected – detected in at least one palmitoylome.

### Palmitoylated Proteins are Overrepresented in PAR-CLIP and Riboprofiling Targets

Given that FASN-bound and translationally regulated RNAs were enriched for transcripts which encode proteins involved in membrane-associated functions, we next asked whether these proteins might also be enriched for known palmitoylation targets. Palmitoylation is a widespread post-translational modification involved in protein trafficking, localization (especially to membranes), and stability^45,46^. We therefore compared the FASN-associated and translationally regulated protein populations with three categories of S-palmitoylation evidence: computationally predicted palmitoylated proteins, proteins identified in large-scale palmitoylome studies, and proteins with experimentally validated palmitoylation sites.

In every category, palmitoylated proteins were statistically overrepresented in both PAR-CLIP targets and riboprofiling hits (Figure 4E). Experimentally validated palmitoylation targets represented 3.2% of PAR-CLIP targets and 3.7% of riboprofiling hits, compared with 2.2% of the global proteome. Taken together with the previously described translational effects, these findings identify an enrichment of proteins associated with palmitoylation among both FASN-associated transcripts and translationally regulated targets. These associations do not establish a mechanistic connection between RNA binding and protein palmitoylation but suggest that post-translationally regulated membrane-associated proteins represent a functional category associated with the FASN RNA and translational landscapes.

## DISCUSSION

FASN has traditionally been studied primarily as a metabolic enzyme, yet increasing evidence has implicated the protein in cellular processes that are not readily explained by its canonical role in fatty acid synthesis^17,27,32^. This study establishes FASN as a broadly interacting cellular RNA-binding protein and defines its RNA-binding landscape in the absence of an exogenous stimulus. We identify RNA interactions involving multiple FASN domains, map thousands of reproducible FASN-associated binding sites across two cell types and find a pronounced enrichment of binding within the 5’ UTRs of mature mRNAs. We further show that FASN-associated transcripts are preferentially represented among transcripts that undergo reduced translational efficiency following FASN depletion. Together, these findings support a role for FASN as an RNA-associated post-transcriptional regulator.

Among mature mRNAs, the pronounced enrichment of FASN binding within the 5’ UTRs is particularly notable given the central role of this region in translational regulation^47^. Previous transcriptome-wide CLIP datasets identified approximately 2% of all binding sites within 5’ UTRs, representing less than 5% of sites mapping to mature mRNAs, compared with approximately 60% and 35% within 3’ UTRs and coding sequences, respectively^34^. This distribution is consistent with the preferential association of several well-characterized RBPs to these regions^48^, including ELAVL1^49,50^, TIAL1^51^, and PUM proteins^52^. Moreover, 3’ UTRs are typically considerably longer than 5’ UTRs, providing more opportunities for RNA-protein interactions^53^. Against this background, the preferential association of FASN with 5’ UTRs is distinctive and provides a plausible connection to its previously reported involvement in translational regulation during viral infection and oncogenic proliferation. Our findings extend these observations by identifying a population of cellular FASN-associated transcripts that exhibit reduced translational efficiency following FASN depletion under basal conditions.

The translational consequences of FASN association depend on where within the transcript binding occurs. Transcripts bound by FASN showed reduced translational efficiency following its depletion, consistent with a positive role for FASN association in their translation. Transcripts bound exclusively within the coding sequence instead showed increased translational efficiency, indicating that FASN association can have opposing consequences at different transcript positions. One interpretation is that FASN influences translation differently depending on the stage of the translational cycle at which it is encountered, with association near the 5’ end affecting initiation or scanning and association within the coding sequence interacting with elongation. This remains speculative, and the mechanism underlying the relationship between FASN binding and translational efficiency is unresolved. The enrichment of palmitoylation-associated proteins among both FASN RNA targets and translationally regulated proteins provides an additional link between the RNA-associated functions of FASN and its established role in lipid metabolism. However, these observations remain correlative and do not establish that FASN RNA binding directly controls palmitoylation, or that palmitoylation mediates the translational effects observed here.

The RNA interactions identified here may also be influenced by cellular context. FASN localization and spatial organization can change in response to metabolic and pathological conditions^54–57^ and such redistribution could alter the population of RNAs encountered by FASN. Conversely, RNA association may itself contribute to the spatial or molecular organization of FASN, positioning it within or alongside complexes that influence the translation of particular transcript populations. Either relationship provides a framework for understanding how a broadly associated metabolic enzyme might acquire distinct regulatory functions in different cellular states, including infection and during oncogenesis^26,27,29,32,55,57^.

## Limitations of the study

This study defines the RNA-binding repertoire of FASN and establishes an association between RNA binding and translational regulation, while several limitations define the scope of these conclusions. PAR-CLIP identifies crosslinkable RNA contacts and therefore does not capture every RNA interaction of FASN. In addition, FASN depletion by siRNA establishes a relationship between FASN abundance and translational efficiency but does not distinguish direct effects of individual RNA interactions from secondary consequences of FASN loss. The enrichment of palmitoylation-associated proteins likewise represents an association rather than evidence that RNA binding controls protein palmitoylation. Finally, this study focuses on the RNA-associated functions of FASN and does not address how these interactions intersect mechanistically with its canonical catalytic activity.

## RESOURCE AVAILABILITY

### Lead contact

Further information and requests for resources and reagents should be directed to and will be fulfilled by the lead contact, Manuel Ascano.

### Data and code availability

All sequencing data are deposited to National Center for Biotechnology Information via Sequence Read Archive (SRA); BioProject ID: *in progress*. All code used for sequencing analysis and figure generation is accessible at https://github.com/Ascano-Lab.

## ACKNOWLEDGMENTS

We thank the Vanderbilt Technologies for Advanced Genomics (VANTAGE) core for sequencing, particularly Angela Jones for assistance in library construction. We thank members of the Ascano laboratory for their support, collegiality, and critical review of the manuscript. This work was supported by the NIH (5R35GM119569).

## AUTHOR CONTRIBUTIONS

Conceptualization, Y.P-P. and M.A.; methodology, Y.P-P., N.M., S.M. and M.A.; Investigation, Y.P-P., J.M., M.S. and S.M.; writing—original draft, Y.P-P.; writing—review & editing, Y.P-P., S.C., S.L. and M.A.; funding acquisition, M.A; resources, N.M., S.M. and M.A.; supervision, M.A.

## DECLARATION OF GENERATIVE AI AND AI-ASSISTED TECHNOLOGIES

Portions of R code and shell scripts used for data parsing and visualization were developed with trouble-shooting assistance from Claude (Claude Sonnet 5.5, Anthropic, 2026) and Google (AI assistant, 2026). All code was reviewed, tested, validated, then reincorporated into author-generated code. The authors take full responsibility for its content.

## SUPPLEMENTAL INFORMATION

**Supplemental Table 1. Statistical analyses appended to figures**.

**Supplemental Table 2. FASN binding sites identified via PAR-CLIP**.

**Supplemental Table 3. Transcripts identified via riboprofiling**.

## METHODS

### Cell lines and culture

U2-OS cell lines were cultured in McCoy’s 5A Medium supplemented to contain 10% fetal bovine serum (FBS), 2 mM L-glutamine, 100 U/mL penicillin, and 100 μg/ml streptomycin. HEK-293T cell lines were cultured in DMEM supplemented to contain 10% FBS and 2 mM L-glutamine. Cells with selection markers were treated with the appropriate antibiotic to its respective concentration: hygromycin (100 μg/ml), blasticidin (25 μg/ml). When necessary, induction of the rtTA expression system was performed by supplementing media to 1 μg/ml of doxycycline. Cell lines were constructed previously by Kim, Arcos, *et al*.^27^.

### Immunoblotting

Samples were separated by SDS-PAGE, proteins semi-dry transferred onto polyvinylidene difluoride membrane, blocked in non-fat dry milk then probed with primary antibody specific to proteins of interest and secondary HRP-conjugated antibodies. Membranes were further processed and analyzed with enhanced chemiluminescent detection.

### PAR-CLIP

PAR-CLIP was performed largely as previously described^58,59^ with the addition of a relabeling step after proteinase K digestion. Briefly, U2-OS and HEK293 cells were grown to 80% confluency (approximately 800 million cells total), then simultaneously doxycycline induced and labeled with 100 µM of 4SU for 16 hours prior to washing, UV_365nm_ irradiation and harvesting. Cells were lysed in NP-40 lysis buffer (50 mM HEPES pH 7.5, 150 mM KCl, 2 mM EDTA, 1 mM NaF, 2% (v/v) NP-40, 0.5 mM DTT, Roche EDTA-free protease inhibitor) and incubated with DynaProtein G beads conjugated with anti-FLAG M2 antibody for 2 hours at 4°C. Beads were washed with high-salt buffer, then samples underwent radiolabeling with γ-^32^P-ATP via CIP and T4 PNK treatment. Protein-RNA complexes were then resolved on a pre-cast 4-20% Bis-Tris NuPage Gradient Gel and regions corresponding to FASN’s molecular weight of 272 kDa were excised. Complexes were electroeluted out of the gel, treated with proteinase K. Liberated RNA was then directly relabeled with T4 PNK (to address possible steric hindrance from FASN’s large size^60,61^). RNA was run on a 15% denaturing urea-PAGE gel and size selected, then adapters were ligated onto both the 3’ and 5’-ends. Adapted RNA was reverse transcribed, and used to generate cDNA libraries, which were then sequenced on the NovaSeq X Series PE150 platform. T-to-C fractions were determined as follows:

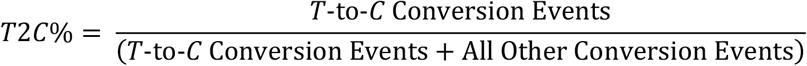

### Binding Site, Motif Determination

PARalyzer detects binding sites through calculation of T-to-C conversion events (2 or 3 events per analyzed transcript) fractions. Kernel density estimates were performed on transcript groups containing at least 5 unique reads in order to determine enrichment of T-to-C converted reads. Binding motifs were analyzed in XSTREME^62^, limiting detection to 3-10 nucleotides and other parameter defaults.

### Knockdown Validation

Knockdowns were validated via RT-qPCR and immunoblotting to detect decreases in RNA and protein levels in surveyed samples.

### RT-qPCR

RNA was collected and extracted with TRIzol. After determination of total RNA concentration via NanoDrop, equal amounts of RNA under each surveyed conditions were reverse transcribed with Super-Script III with oligo d(T) primer. Real-time PCR reactions were performed using Fast SYBR Green on a QuantStudio 3 machine. Obtained target Ct values were normalized to *TUBA1A* Ct values and used to calculate ΔCt, with relative expression calculated using the ΔΔCt method.

### Ribo-seq

Ribosome profiling was performed largely as previously described^63^, with minor modifications to optimize the procedure for U2-OS cells. Briefly, each experimental condition was seeded in 4 15 cm plates, 5 million cells per plate. Knockdown samples were seeded four days prior to collection, then treated the next day with 1500 pmol of siRNA transfected with Lipofectamine 2000 (in accordance with the manufacturer’s instructions) for 72 hours of knockdown. Cells overexpressing or expressing baseline levels of FASN were seeded two days prior to collection. All cells were induced with doxycycline for 16 hours prior to collection. All samples were then washed and placed in liquid nitrogen to halt translation. Cells were then immediately lysed on ice in polysome lysis buffer (1% (v/v) Triton X-100, 25 U/mL Turbo DNase I in polysome buffer – 20 mM Tris pH 7.4, 150 mM NaCl, 5 mM MgCl_2_, 1 mM DTT, 100 µg/mL cycloheximide), with solidified lysate scraped and pooled together to thaw on ice. Lysates from experimental conditions were joined together to form a single aliquot, triturated through a 26G needle and clarified of cell debris, and 100 μL (representing approximately 2% of the lysate volume) was reserved directly into 900 μL of TRIzol to serve as an input control. RNA was footprinted to the ribosome with RNase I, then ribosome-RNA complexes were purified on a 1 M sucrose cushion and RNA subsequently isolated. RNA was run on a pre-cast 15% TBE-urea polyacrylamide gel and size selected, then adapters were ligated onto the 3’-end. Adapted RNA was reverse transcribed, and used to generate barcoded cDNA libraries, which were then sequenced on the NovaSeq X Series PE150 platform.

### RNA Sequencing

Input RNA was processed where indicated through TRIzol per the manufacturer’s instructions. RNA was reverse transcribed, and used to generate barcoded cDNA libraries, which were then sequenced on the NovaSeq X Series PE150 platform.

### Translation Efficiency Determination

Translational efficiency was calculated as the ratio of the ribosome protected fragments (RPFs) to input RNA-seq signal and modeled using deltaTE^43^, with differential translation assessed as the interaction between read type and condition. Transcripts were assigned to regulatory classes according to which contrasts reached significance: exclusive (significant change in TE without a corresponding change in RNA), intensified or buffered (changes in TE that reinforce or oppose the RNA change), and forwarded (proportional changes in RNA and RPF, indicating transcript-level regulation without a statistically change in TE). Overrepresentation of lowly expressed transcripts was minimized utilizing approximate posterior estimation for generalized linear models (“apeglm”), as previously described^64^.

### Statistical Analyses

Similarities between cluster distributions were determined using a chi-squared test. Significant differences between cumulative distribution fractions were calculated using a 10,000-permutation bootstrapped Kolmogorov-Smirnov test (with identical seeds across comparisons) and a Mann-Whitney *U* test. Monotonic trends in TE across ordered groups were tested with a two-sided Jonckheere-Terpstra test. In all instances of multiple comparisons within an analysis family, p-values were adjusted by the Bonferroni method. Hypergeometric analysis was utilized to determine overrepresentation of a target protein population compared to global proteomes, with false discovery rates within groups of tests correcting adjusting *p-*values utilizing the Benjamini-Hochberg procedure. Significant threshold of 0.05 was assumed for all statistical tests. Number of biological replicates of assays are defined within the corresponding figure. Error bars show in figures represent means ± standard deviation.

## KEY RESOURCES TABLE

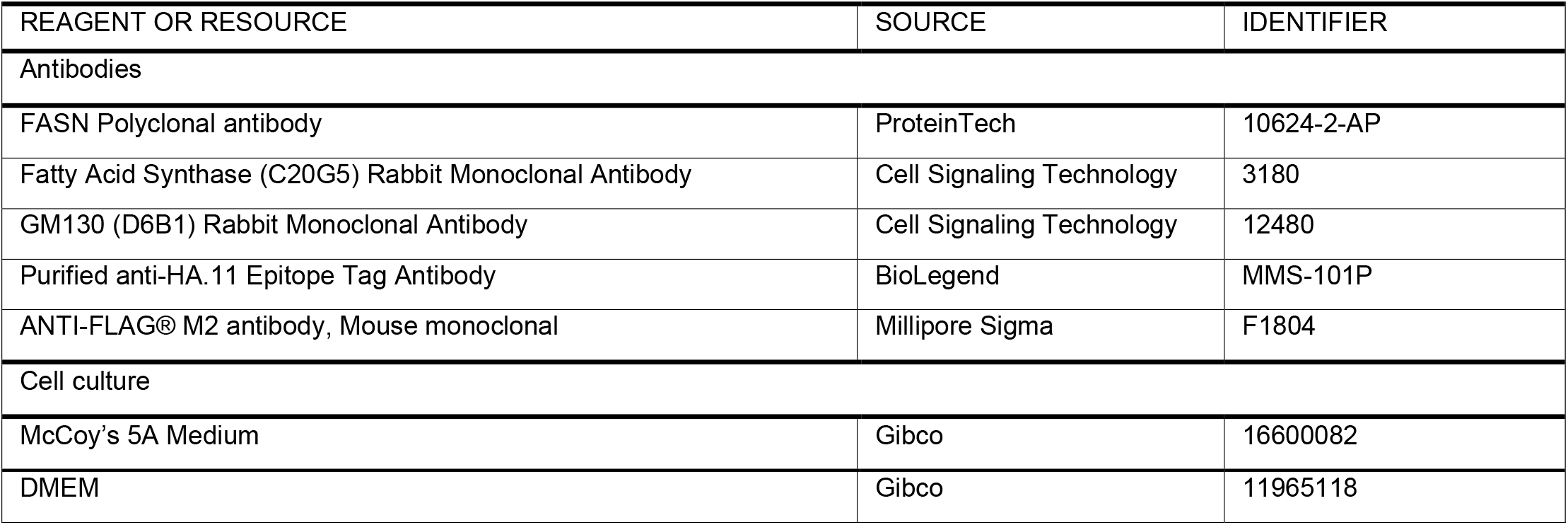
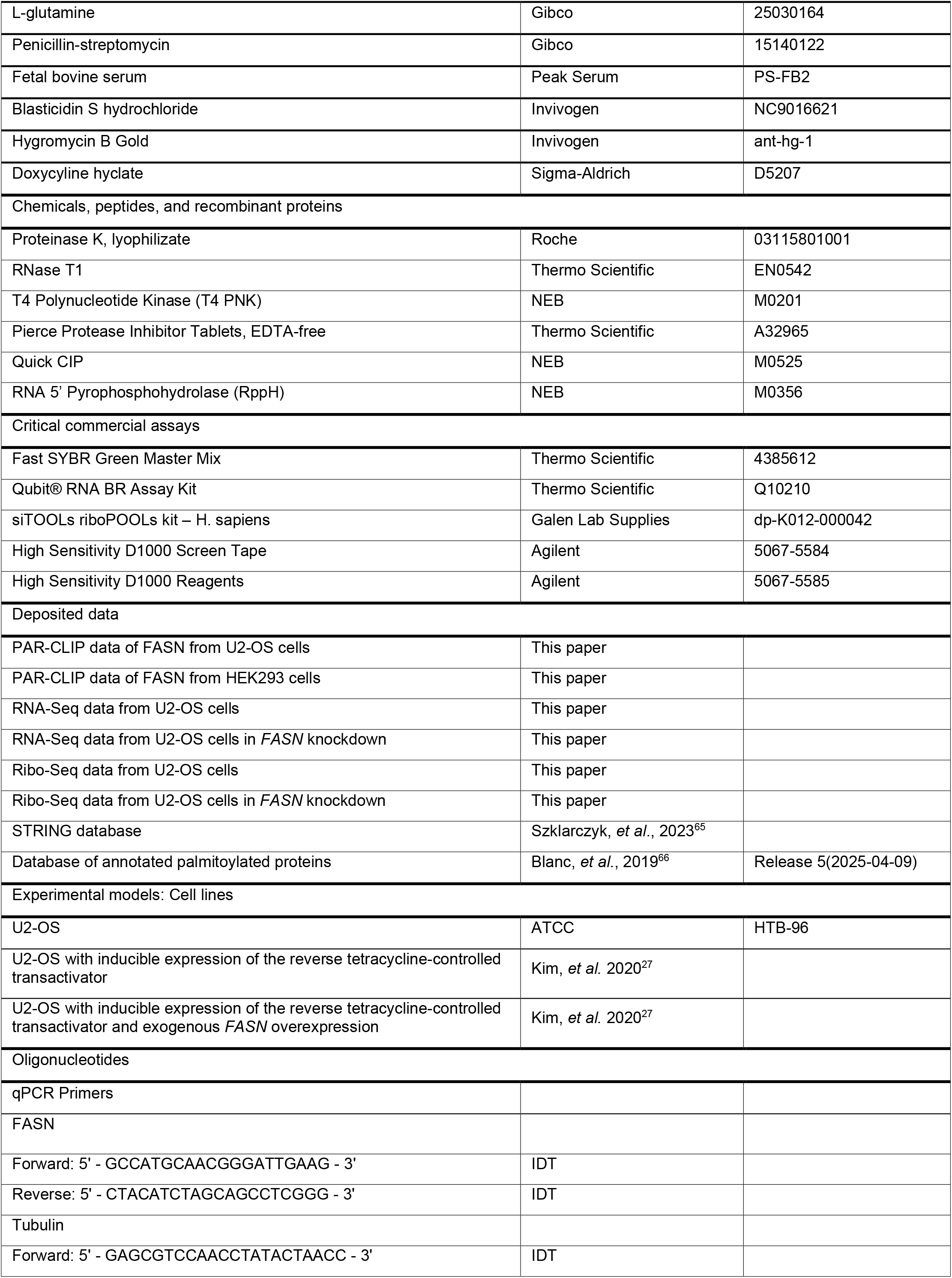
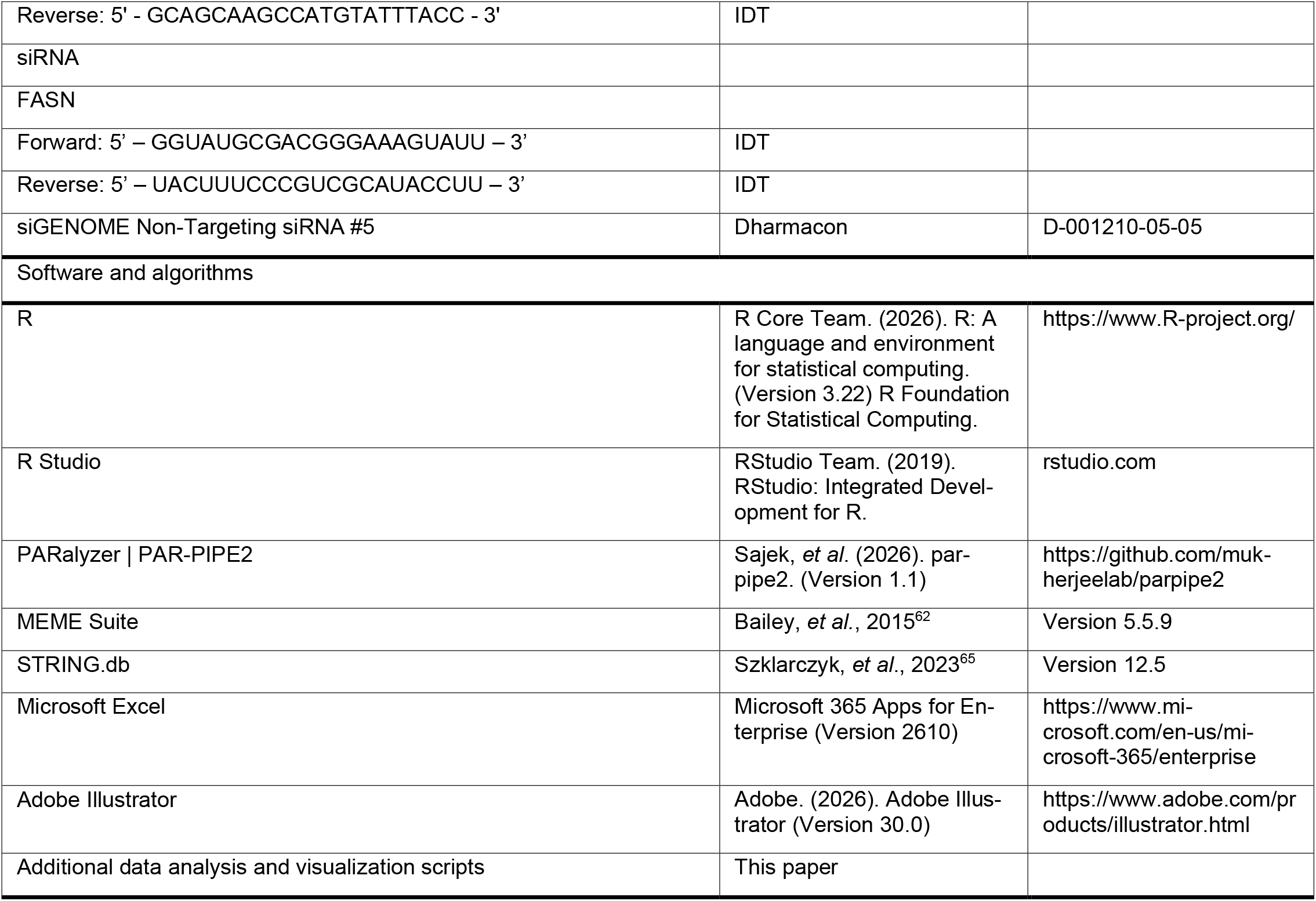

## REFERENCES

1. Castello, A., Fischer, B., Eichelbaum, K., Horos, R., Beckmann, B.M., Strein, C., Davey, N.E., Humphreys, D.T., Preiss, T., Steinmetz, L.M., et al. (2012). Insights into RNA biology from an atlas of mammalian mRNA-binding proteins. Cell 149, 1393–1406. 10.1016/j.cell.2012.04.031.

2. He, C., Sidoli, S., Warneford-Thomson, R., Tatomer, D.C., Wilusz, J.E., Garcia, B.A., and Bonasio, R. (2016). High-Resolution Mapping of RNA-Binding Regions in the Nuclear Proteome of Embryonic Stem Cells. Mol Cell 64, 416–430. 10.1016/j.molcel.2016.09.034.

3. Queiroz, R.M.L., Smith, T., Villanueva, E., Marti-Solano, M., Monti, M., Pizzinga, M., Mirea, D.M., Ramakrishna, M., Harvey, R.F., Dezi, V., et al. (2019). Comprehensive identification of RNA-protein interactions in any organism using orthogonal organic phase separation (OOPS). Nat Biotechnol 37, 169–178. 10.1038/s41587-018-0001-2.

4. Shchepachev, V., Bresson, S., Spanos, C., Petfalski, E., Fischer, L., Rappsilber, J., and Tollervey, D. (2019). Defining the RNA interactome by total RNA-associated protein purification. Mol Syst Biol 15, e8689. 10.15252/msb.20188689.

5. Hentze, M.W., Sommerkamp, P., Ravi, V., and Gebauer, F. (2025). Rethinking RNA-binding proteins: Riboregulation challenges prevailing views. Cell 188, 4811–4827. 10.1016/j.cell.2025.06.021.

6. Sahadevan, S., Sekaran, T., Hentze, M.W., Perez-Perri, J.I., and Schwarzl, T. (2026). Signal or noise? RNA-binding proteins and the challenges of binding site assignments. Nucleic Acids Res 54. 10.1093/nar/gkag452.

7. Huppertz, I., Perez-Perri, J.I., Mantas, P., Sekaran, T., Schwarzl, T., Russo, F., Ferring-Appel, D., Koskova, Z., Dimitrova-Paternoga, L., Kafkia, E., et al. (2022). Riboregulation of Enolase 1 activity controls glycolysis and embryonic stem cell differentiation. Mol Cell 82, 2666–2680 e2611. 10.1016/j.molcel.2022.05.019.

8. Zhang, Q., Cheng, S., Gu, Y., Pan, Y., Zhu, X., Sun, J., Wen, Y., and Gao, S. (2026). P5CS represses tumor progression via inhibiting assembly of 48S pre-translation initiation complex. Dev Cell 61, 1521–1534 e1527. 10.1016/j.devcel.2026.05.011.

9. Chirala, S.S., Jayakumar, A., Gu, Z.W., and Wakil, S.J. (2001). Human fatty acid synthase: role of interdomain in the formation of catalytically active synthase dimer. Proc Natl Acad Sci U S A 98, 3104–3108. 10.1073/pnas.051635998.

10. Ameer, F., Scandiuzzi, L., Hasnain, S., Kalbacher, H., and Zaidi, N. (2014). De novo lipogenesis in health and disease. Metabolism 63, 895–902. 10.1016/j.metabol.2014.04.003.

11. Chorna, N.E., Santos-Soto, I.J., Carballeira, N.M., Morales, J.L., de la Nuez, J., Catala-Valentin, A., Chornyy, A.P., Vazquez-Montes, A., and De Ortiz, S.P. (2013). Fatty acid synthase as a factor required for exercise-induced cognitive enhancement and dentate gyrus cellular proliferation. PLoS One 8, e77845. 10.1371/journal.pone.0077845.

12. Gonzalez-Bohorquez, D., Gallego Lopez, I.M., Jaeger, B.N., Pfammatter, S., Bowers, M., Semenkovich, C.F., and Jessberger, S. (2022). FASN-dependent de novo lipogenesis is required for brain development. Proc Natl Acad Sci U S A 119. 10.1073/pnas.2112040119.

13. Bowers, M., Liang, T., Gonzalez-Bohorquez, D., Zocher, S., Jaeger, B.N., Kovacs, W.J., Rohrl, C., Cramb, K.M.L., Winterer, J., Kruse, M., et al. (2020). FASN-Dependent Lipid Metabolism Links Neurogenic Stem/Progenitor Cell Activity to Learning and Memory Deficits. Cell Stem Cell 27, 98–109 e111. 10.1016/j.stem.2020.04.002.

14. Cuyas, E., Pedarra, S., Verdura, S., Pardo, M.A., Espin Garcia, R., Serrano-Hervas, E., Llop-Hernandez, A., Teixidor, E., Bosch-Barrera, J., Lopez-Bonet, E., et al. (2024). Fatty acid synthase (FASN) is a tumor-cell-intrinsic metabolic checkpoint restricting T-cell immunity. Cell Death Discov 10, 417. 10.1038/s41420-024-02184-z.

15. Drury, J., Young, L.E.A., Scott, T.L., Kelson, C.O., He, D., Liu, J., Wu, Y., Wang, C., Weiss, H.L., Fan, T., et al. (2022). Tissue-Specific Downregulation of Fatty Acid Synthase Suppresses Intestinal Adenoma Formation via Coordinated Reprograming of Transcriptome and Metabolism in the Mouse Model of Apc-Driven Colorectal Cancer. Int J Mol Sci 23. 10.3390/ijms23126510.

16. Scaglia, N., Tyekucheva, S., Zadra, G., Photopoulos, C., and Loda, M. (2014). De novo fatty acid synthesis at the mitotic exit is required to complete cellular division. Cell Cycle 13, 859–868. 10.4161/cc.27767.

17. Pasquier, C., Messmer, M., Moroge, L., Knol, L., Chicher, J., Ngondo, R.P., Pfeffer, S., and Girardi, E. (2026). Fatty Acid Synthase binds endogenous dsRNAs and dampens the innate immune response to exogenous dsRNAs by limiting their accumulation. bioRxiv. 10.1101/2025.07.16.662511.

18. Wang, K., Li, L., Jin, J., An, Y., Wang, Z., Zhou, S., Zhang, J., Abuduaini, B., Cheng, C., and Li, N. (2022). Fatty acid synthase (Fasn) inhibits the expression levels of immune response genes via alteration of alternative splicing in islet cells. J Diabetes Complications 36, 108159. 10.1016/j.jdiacomp.2022.108159.

19. Wei, H., Weaver, Y.M., Yang, C., Zhang, Y., Hu, G., Karner, C.M., Sieber, M., DeBerardinis, R.J., and Weaver, B.P. (2024). Proteolytic activation of fatty acid synthase signals pan-stress resolution. Nat Metab 6, 113–126. 10.1038/s42255-023-00939-z.

20. Nuzzo, P.V., Rodrigues, S., Ribeiro, C.F., Teixeira, I.F., Fanelli, G.N., Bleve, S., Ravera, F., Pakula, H., Pederzoli, F., Nanus, D.M., and Loda, M. (2025). Targeting cancer metabolism: Therapeutic potential of the fatty acid synthase (FASN) inhibitors. Crit Rev Oncol Hematol 214, 104910. 10.1016/j.critrevonc.2025.104910.

21. Fhu, C.W., and Ali, A. (2020). Fatty Acid Synthase: An Emerging Target in Cancer. Molecules 25. 10.3390/molecules25173935.

22. Chen, L.Y., Wu, D.S., and Shen, Y.A. (2024). Fatty acid synthase inhibitor cerulenin hinders liver cancer stem cell properties through FASN/APP axis as novel therapeutic strategies. J Lipid Res 65, 100660. 10.1016/j.jlr.2024.100660.

23. Banerjee, M., Zaytseva, Y.Y., Reusch, E.M., Napier, D.L., Hasani, S., Rychahou, P., Izumi, T., Cheek, D.A., Li, J., Flight, R.M., et al. (2026). FASN Inhibition Enhances the Efficacy of Chemotherapy in Colorectal Cancer by Inhibiting the DNA Damage Response. Cancer Res 86, 2522–2536. 10.1158/0008-5472.CAN-25-1917.

24. Huang, Y., Xu, Q., Zheng, L., Zheng, H., Chen, H., Li, Y., and Ma, S. (2026). The implications of FASN in viral infection and related diseases: a promising target in antiviral therapies. Front Cell Infect Microbiol 16, 1846176. 10.3389/fcimb.2026.1846176.

25. Fernandez-Chamorro, J., Francisco-Velilla, R., Ramajo, J., and Martinez-Salas, E. (2019). Rab1b and ARF5 are novel RNA-binding proteins involved in FMDV IRES-driven RNA localization. Life Sci Alliance 2. 10.26508/lsa.201800131.

26. Liu, Y.Y., Liang, X.D., Liu, C.C., Cheng, Y., Chen, H., Baloch, A.S., Zhang, J., Go, Y.Y., and Zhou, B. (2021). Fatty Acid Synthase Is Involved in Classical Swine Fever Virus Replication by Interaction with NS4B. J Virol 95, e0078121. 10.1128/JVI.00781-21.

27. Kim, B., Arcos, S., Rothamel, K., Jian, J., Rose, K.L., McDonald, W.H., Bian, Y., Reasoner, S., Barrows, N.J., Bradrick, S., et al. (2020). Discovery of Widespread Host Protein Interactions with the Pre-replicated Genome of CHIKV Using VIR-CLASP. Mol Cell 78, 624–640 e627. 10.1016/j.molcel.2020.04.013.

28. Flynn, R.A., Belk, J.A., Qi, Y., Yasumoto, Y., Wei, J., Alfajaro, M.M., Shi, Q., Mumbach, M.R., Limaye, A., DeWeirdt, P.C., et al. (2021). Discovery and functional interrogation of SARS-CoV-2 RNA-host protein interactions. Cell 184, 2394–2411 e2316. 10.1016/j.cell.2021.03.012.

29. Heaton, N.S., Perera, R., Berger, K.L., Khadka, S., Lacount, D.J., Kuhn, R.J., and Randall, G. (2010). Dengue virus nonstructural protein 3 redistributes fatty acid synthase to sites of viral replication and increases cellular fatty acid synthesis. Proc Natl Acad Sci U S A 107, 17345–17350. 10.1073/pnas.1010811107.

30. Albano, C., Trifiro, L., Hewelt-Belka, W., Cairns, D.M., Pasquero, S., Griffante, G., Gugliesi, F., Bajetto, G., Garwolinska, D., Rossi, M., et al. (2025). The impact of fatty acid synthase on HSV-1 infection dynamics. PLoS Pathog 21, e1013068. 10.1371/journal.ppat.1013068.

31. Huang, J.T., Tseng, C.P., Liao, M.H., Lu, S.C., Yeh, W.Z., Sakamoto, N., Chen, C.M., and Cheng, J.C. (2013). Hepatitis C virus replication is modulated by the interaction of nonstructural protein NS5B and fatty acid synthase. J Virol 87, 4994–5004. 10.1128/JVI.02526-12.

32. Kapadia, B., Nanaji, N.M., Bhalla, K., Bhandary, B., Lapidus, R., Beheshti, A., Evens, A.M., and Gartenhaus, R.B. (2018). Fatty Acid Synthase induced S6Kinase facilitates USP11-eIF4B complex formation for sustained oncogenic translation in DLBCL. Nat Commun 9, 829. 10.1038/s41467-018-03028-y.

33. Castello, A., Fischer, B., Frese, C.K., Horos, R., Alleaume, A.M., Foehr, S., Curk, T., Krijgsveld, J., and Hentze, M.W. (2016). Comprehensive Identification of RNA-Binding Domains in Human Cells. Mol Cell 63, 696–710. 10.1016/j.molcel.2016.06.029.

34. Yang, Y.C., Di, C., Hu, B., Zhou, M., Liu, Y., Song, N., Li, Y., Umetsu, J., and Lu, Z.J. (2015). CLIPdb: a CLIP-seq database for protein-RNA interactions. BMC Genomics 16, 51. 10.1186/s12864-015-1273-2.

35. Mullari, M., Lyon, D., Jensen, L.J., and Nielsen, M.L. (2017). Specifying RNA-Binding Regions in Proteins by Peptide Cross-Linking and Affinity Purification. J Proteome Res 16, 2762–2772. 10.1021/acs.jproteome.7b00042.

36. Trendel, J., Schwarzl, T., Horos, R., Prakash, A., Bateman, A., Hentze, M.W., and Krijgsveld, J. (2019). The Human RNA-Binding Proteome and Its Dynamics during Translational Arrest. Cell 176, 391–403 e319. 10.1016/j.cell.2018.11.004.

37. Baltz, A.G., Munschauer, M., Schwanhausser, B., Vasile, A., Murakawa, Y., Schueler, M., Youngs, N., Penfold-Brown, D., Drew, K., Milek, M., et al. (2012). The mRNA-bound proteome and its global occupancy profile on protein-coding transcripts. Mol Cell 46, 674–690. 10.1016/j.molcel.2012.05.021.

38. Urdaneta, E.C., Vieira-Vieira, C.H., Hick, T., Wessels, H.H., Figini, D., Moschall, R., Medenbach, J., Ohler, U., Granneman, S., Selbach, M., and Beckmann, B.M. (2019). Purification of cross-linked RNA-protein complexes by phenol-toluol extraction. Nat Commun 10, 990. 10.1038/s41467-019-08942-3.

39. Qin, W., Myers, S.A., Carey, D.K., Carr, S.A., and Ting, A.Y. (2021). Spatiotemporally-resolved mapping of RNA binding proteins via functional proximity labeling reveals a mitochondrial mRNA anchor promoting stress recovery. Nat Commun 12, 4980. 10.1038/s41467-021-25259-2.

40. Harris, M.E., and Christian, E.L. (2009). RNA crosslinking methods. Methods Enzymol 468, 127–146. 10.1016/S0076-6879(09)68007-1.

41. Armaos, A., Colantoni, A., Proietti, G., Rupert, J., and Tartaglia, G.G. (2021). catRAPID omics v2.0: going deeper and wider in the prediction of protein-RNA interactions. Nucleic Acids Res 49, W72–W79. 10.1093/nar/gkab393.

42. Chirala, S.S., Chang, H., Matzuk, M., Abu-Elheiga, L., Mao, J., Mahon, K., Finegold, M., and Wakil, S.J. (2003). Fatty acid synthesis is essential in embryonic development: fatty acid synthase null mutants and most of the heterozygotes die in utero. Proc Natl Acad Sci U S A 100, 6358–6363. 10.1073/pnas.0931394100.

43. Chothani, S., Adami, E., Ouyang, J.F., Viswanathan, S., Hubner, N., Cook, S.A., Schafer, S., and Rackham, O.J.L. (2019). deltaTE: Detection of Translationally Regulated Genes by Integrative Analysis of Ribo-seq and RNA-seq Data. Curr Protoc Mol Biol 129, e108. 10.1002/cpmb.108.

44. Knowles, L.M., Axelrod, F., Browne, C.D., and Smith, J.W. (2004). A fatty acid synthase blockade induces tumor cell-cycle arrest by down-regulating Skp2. J Biol Chem 279, 30540–30545. 10.1074/jbc.M405061200.

45. Linder, M.E., and Deschenes, R.J. (2007). Palmitoylation: policing protein stability and traffic. Nat Rev Mol Cell Biol 8, 74–84. 10.1038/nrm2084.

46. Salaun, C., Greaves, J., and Chamberlain, L.H. (2010). The intracellular dynamic of protein palmitoylation. J Cell Biol 191, 1229–1238. 10.1083/jcb.201008160.

47. Leppek, K., Das, R., and Barna, M. (2018). Functional 5’ UTR mRNA structures in eukaryotic translation regulation and how to find them. Nat Rev Mol Cell Biol 19, 158–174. 10.1038/nrm.2017.103.

48. Grzybowska, E.A., and Wakula, M. (2021). Protein Binding to Cis-Motifs in mRNAs Coding Sequence Is Common and Regulates Transcript Stability and the Rate of Translation. Cells 10. 10.3390/cells10112910.

49. Lebedeva, S., Jens, M., Theil, K., Schwanhausser, B., Selbach, M., Landthaler, M., and Rajewsky, N. (2011). Transcriptome-wide analysis of regulatory interactions of the RNA-binding protein HuR. Mol Cell 43, 340–352. 10.1016/j.molcel.2011.06.008.

50. Rothamel, K., Arcos, S., Kim, B., Reasoner, C., Lisy, S., Mukherjee, N., and Ascano, M. (2021). ELAVL1 primarily couples mRNA stability with the 3’ UTRs of interferon-stimulated genes. Cell Rep 35, 109178. 10.1016/j.celrep.2021.109178.

51. Meyer, C., Garzia, A., Mazzola, M., Gerstberger, S., Molina, H., and Tuschl, T. (2018). The TIA1 RNA-Binding Protein Family Regulates EIF2AK2-Mediated Stress Response and Cell Cycle Progression. Mol Cell 69, 622–635 e626. 10.1016/j.molcel.2018.01.011.

52. Sternburg, E.L., Lillibridge, J.J., Phandthong, R., and Karginov, F.V. (2023). Mammalian pumilio proteins control cellular morphology, migration, and adhesion. Sci Rep 13, 3002. 10.1038/s41598-023-30004-4.

53. Mignone, F., Gissi, C., Liuni, S., and Pesole, G. (2002). Untranslated regions of mRNAs. Genome Biol 3, REVIEWS0004. 10.1186/gb-2002-3-3-reviews0004.

54. Suresh, H.G., da Silveira Dos Santos, A.X., Kukulski, W., Tyedmers, J., Riezman, H., Bukau, B., and Mogk, A. (2015). Prolonged starvation drives reversible sequestration of lipid biosynthetic enzymes and organelle reorganization in Saccharomyces cerevisiae. Mol Biol Cell 26, 1601–1615. 10.1091/mbc.E14-11-1559.

55. Madigan, A.A., Rycyna, K.J., Parwani, A.V., Datiri, Y.J., Basudan, A.M., Sobek, K.M., Cummings, J.L., Basse, P.H., Bacich, D.J., and O’Keefe, D.S. (2014). Novel nuclear localization of fatty acid synthase correlates with prostate cancer aggressiveness. Am J Pathol 184, 2156–2162. 10.1016/j.ajpath.2014.04.012.

56. Di Vizio, D., Adam, R.M., Kim, J., Kim, R., Sotgia, F., Williams, T., Demichelis, F., Solomon, K.R., Loda, M., Rubin, M.A., et al. (2008). Caveolin-1 interacts with a lipid raft-associated population of fatty acid synthase. Cell Cycle 7, 2257–2267. 10.4161/cc.7.14.6475.

57. Ma, X., Gu, W., Li, X., Ding, X., and Zhu, L. (2025). Fatty acid synthase may facilitate the trafficking of bovine alpha herpesvirus 1 out of the Golgi apparatus, potentially promoting viral infection. Microbiol Spectr 13, e0138825. 10.1128/spectrum.01388-25.

58. Lisy, S., Rothamel, K., Perevalova-Pinzul, Y., and Ascano, M. (2024). PAR-dCLIP: Enabling detection of RNA binding protein target transcripts bound at 5’ termini through the incorporation of a decapping step. Methods Enzymol 705, 159–222. 10.1016/bs.mie.2024.08.003.

59. Spitzer, J., Hafner, M., Landthaler, M., Ascano, M., Farazi, T., Wardle, G., Nusbaum, J., Khorshid, M., Burger, L., Zavolan, M., and Tuschl, T. (2014). PAR-CLIP (Photoactivatable Ribonucleoside-Enhanced Crosslinking and Immunoprecipitation): a step-by-step protocol to the transcriptome-wide identification of binding sites of RNA-binding proteins. Methods Enzymol 539, 113–161. 10.1016/B978-0-12-420120-0.00008-6.

60. Maier, T., Jenni, S., and Ban, N. (2006). Architecture of mammalian fatty acid synthase at 4.5 A resolution. Science 311, 1258–1262. 10.1126/science.1123248.

61. Maier, T., Leibundgut, M., and Ban, N. (2008). The crystal structure of a mammalian fatty acid synthase. Science 321, 1315–1322. 10.1126/science.1161269.

62. Bailey, T.L., Johnson, J., Grant, C.E., and Noble, W.S. (2015). The MEME Suite. Nucleic Acids Res 43, W39–49. 10.1093/nar/gkv416.

63. McGlincy, N.J., and Ingolia, N.T. (2017). Transcriptome-wide measurement of translation by ribosome profiling. Methods 126, 112–129. 10.1016/j.ymeth.2017.05.028.

64. Zhu, A., Ibrahim, J.G., and Love, M.I. (2019). Heavy-tailed prior distributions for sequence count data: removing the noise and preserving large differences. Bioinformatics 35, 2084–2092. 10.1093/bioinformatics/bty895.

65. Szklarczyk, D., Kirsch, R., Koutrouli, M., Nastou, K., Mehryary, F., Hachilif, R., Gable, A.L., Fang, T., Doncheva, N.T., Pyysalo, S., et al. (2023). The STRING database in 2023: protein-protein association networks and functional enrichment analyses for any sequenced genome of interest. Nucleic Acids Res 51, D638–D646. 10.1093/nar/gkac1000.

66. Blanc, M., David, F.P.A., and van der Goot, F.G. (2019). SwissPalm 2: Protein S-Palmitoylation Database. Methods Mol Biol 2009, 203–214. 10.1007/978-1-4939-9532-5_16.

