## Supplemental Figures for "Fatty Acid Synthase Binds Cellular mRNAs and its Depletion Reduces Translational Efficiency"

#### Supplemental Legends

**Figure S1. Fatty Acid Synthase contains multiple RNA binding domains which bind to a range of RNA subpopulations.**

(A) PAR-CLIP precipitation of overexpressed FASN in U2-OS.
(B) $^{32}$P labeled-RNA isolated after immunoprecipitation of overexpressed FASN.

**Figure S2. Fatty Acid Synthase interacts with a wide range of Pol-II mediated transcripts.**

(A) Correlation of annotated FASN binding site T-to-C enrichment identified across biological replicates in U2-OS cells.
(B) Correlation of annotated FASN binding site T-to-C enrichment identified across biological replicates in HEK cells.
(C) Unionized analysis of all binding sites identified in a global analysis of HEK and U2-OS PAR-CLIP libraries, including raw cluster count and kernel density estimation (KDE) that a cluster will be identified with a given T-to-C enrichment score.
(D) Binding sites identified via PAR-CLIP when unionizing HEK and U2-OS libraires.
(E) Binding site distribution across different levels of T-to-C conversion enrichment values in U2-OS cells (*n* = 2 biological replicates).
(F) Overlap of transcripts with an assigned ENSEMBL ID identified as containing at least one binding site (*n* = 2 biological replicates per line).
(G) Overlap of annotated gene RNA binding sites as a result of individual or unified site detection (*n* = 2 biological replicates per line).
(H) Distribution of clusters in mRNA regions identified via PAR-CLIP.
(I) Cumulative distribution analysis of T-to-C transitions in PAR-CLIP targets mapping to mRNA.
(J) Gene ontology analysis of protein-coding hits, querying localization to cellular compartments.
(K) Gene ontology analysis of protein-coding hits, querying molecular function.
(L) Gene ontology analysis of protein-coding hits, querying biological processes.
(M) Top RNA binding motifs identified.

**Figure S3. Effects of Fatty Acid Synthase on translation.**

(A) Replicate mean of 99-precentile winsorized RPF RPKM to input RNA RPKM in in cells undergoing FASN knockdown or treatment with a non-targeting control in U2-OS, mapped along a normalized gene body (*n* = 2 per condition).
(B) Protein-coding transcripts with statistically significant differences in RNA expression, RPF enrichment, and/or translational efficiencies in cells undergoing FASN knockdown in U2-OS (*n* = 2 per condition).
(C) Cumulative distribution analysis of translational efficiencies of statistically significant protein coding-transcripts (as in S3B) targets bound and unbound by FASN in FASN knockdown conditions.
Inset: Overlap of annotated transcripts identified as statistically significant riboprofiling hits
(D) Cumulative distribution analysis of translational efficiencies of protein-coding transcripts categorized as “forwarded” (as in S3B) targets bound and unbound by FASN in FASN knockdown conditions.
(E) Box-and-whisker plot of TE $\log_2$ fold changes of statistically significant protein coding-transcripts (as in S3B). Lines within boxes represent medians.
(F) Cumulative distribution analysis of translational efficiencies of mRNA targets with a singular FASN binding site in the CDS, opposing all other categories with a single binding site. Categories are gene-exclusive.
(G) Cumulative distribution analysis of translational efficiencies of mRNA targets by quartile of T-to-C enrichment of the most enriched binding site. Genes at the bin cutoff were randomly assigned in cases of ties. Approximate maximum T-to-C conversion fold enrichments are 10 (bottom 25%), 18 (25-50%), 36 (50-75%).

(H) Pol-II transcripts with statistically significant differences in RNA expression, RPF enrichment, and/or translational efficiencies in cells undergoing FASN overexpression in U2-OS ($n$ = 2 per condition).

* - statistical significance (refer to Table S1)

**Figure S4. Ontological analysis and palmitoylation status of FASN-regulated genes.**

(A) Gene ontology analysis of protein-coding hits with negative TEs, querying molecular function.

(B) Gene ontology analysis of protein-coding hits with negative TEs and containing at least one FASN binding site, querying molecular function.

(C) Gene ontology analysis of protein-coding hits with negative TEs and containing at least one FASN binding site at the 5' UTR, querying cellular compartment.

**Supplemental Figure 1.**

**A**

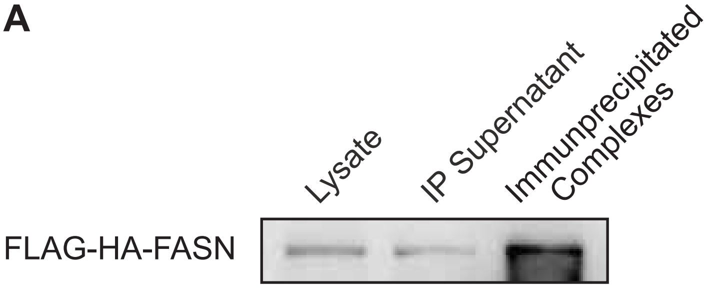


**B**

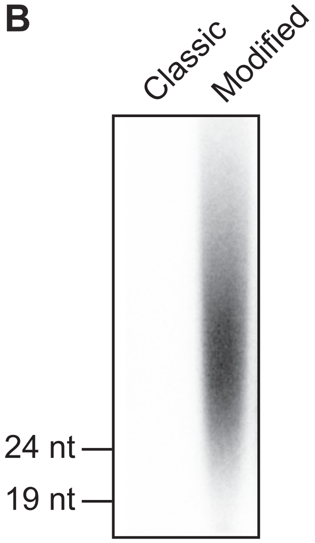

#### Supplemental Figure 2.

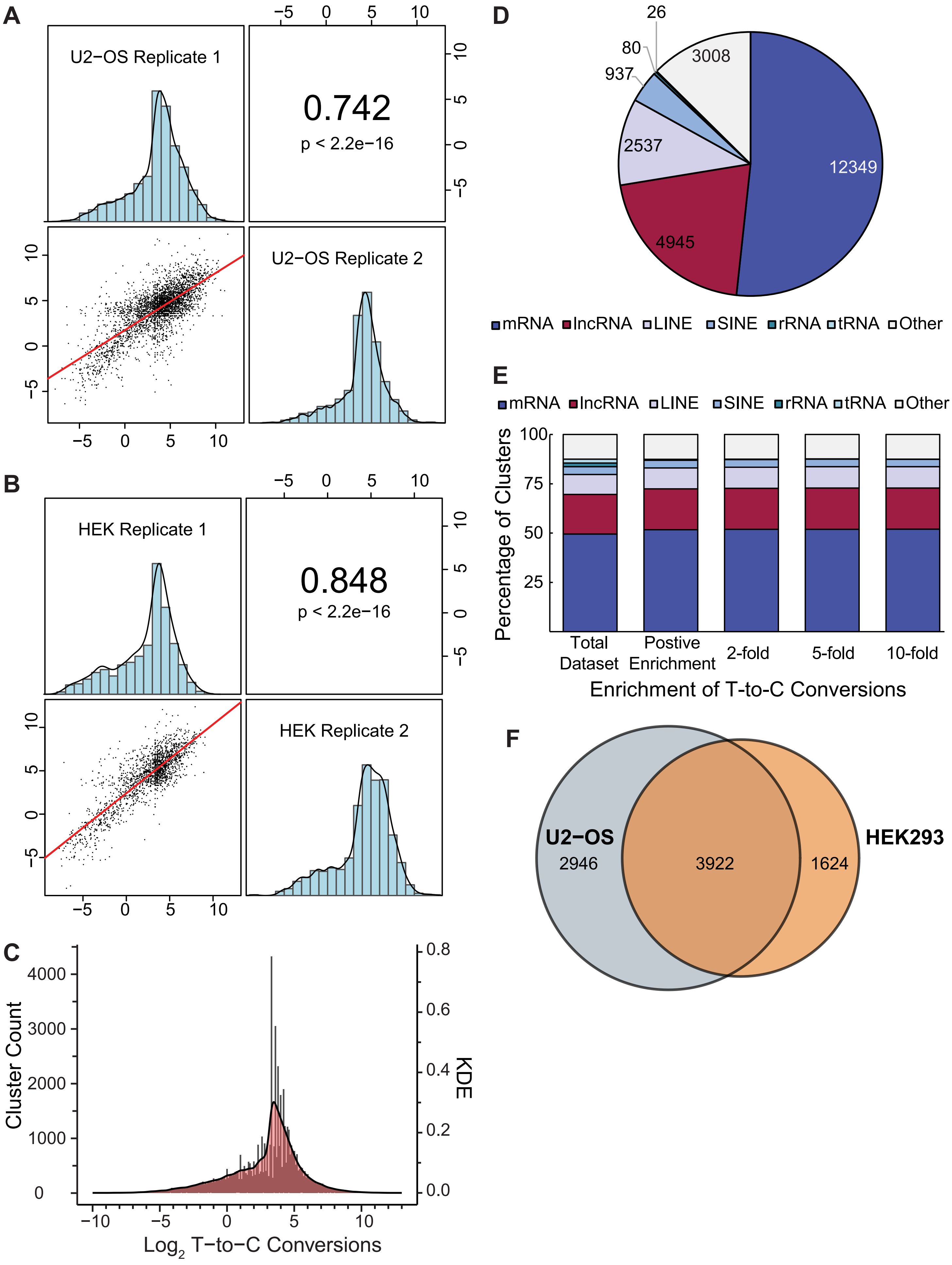

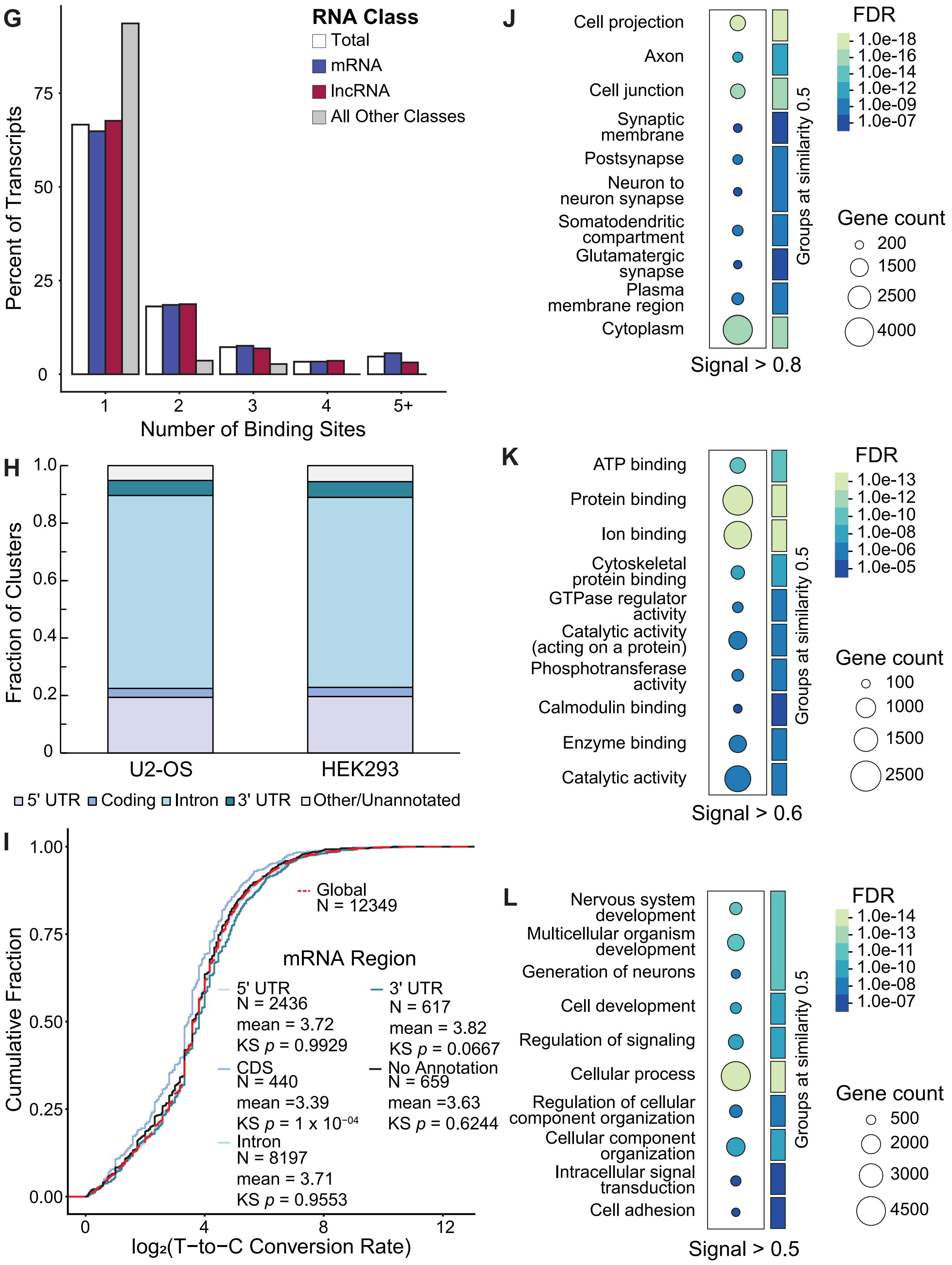
G
RNA Class
Total
mRNA
lncRNA
All Other Classes
Percent of Transcripts
0
25
50
75
1
2
3
4
5+
Number of Binding Sites
H
Fraction of Clusters
0
0.2
0.4
0.6
0.8
1.0
U2-OS
HEK293
5' UTR
Coding
Intron
3' UTR
Other/Unannotated
I
Cumulative Fraction
0.00
0.25
0.50
0.75
1.00
0
4
8
12
log2(T−to−C Conversion Rate)
Global
N = 12349
mRNA Region
5' UTR
N = 2436
mean = 3.72
KS p = 0.9929
CDS
N = 440
mean =3.39
KS p = 1 x 10−04
Intron
N = 8197
mean = 3.71
KS p = 0.9553
3' UTR
N = 617
mean = 3.82
KS p = 0.0667
No Annotation
N = 659
mean =3.63
KS p = 0.6244
J
Cell projection
Axon
Cell junction
Synaptic membrane
Postsynapse
Neuron to neuron synapse
Somatodendritic compartment
Glutamatergic synapse
Plasma membrane region
Cytoplasm
Groups at similarity 0.5
Signal > 0.8
FDR
1.0e-18
1.0e-16
1.0e-14
1.0e-12
1.0e-09
1.0e-07
Gene count
200
1500
2500
4000
K
ATP binding
Protein binding
Ion binding
Cytoskeletal protein binding
GTPase regulator activity
Catalytic activity (acting on a protein)
Phosphotransferase activity
Calmodulin binding
Enzyme binding
Catalytic activity
Groups at similarity 0.5
Signal > 0.6
FDR
1.0e-13
1.0e-12
1.0e-10
1.0e-08
1.0e-06
1.0e-05
Gene count
100
1000
1500
2500
L
Nervous system development
Multicellular organism development
Generation of neurons
Cell development
Regulation of signaling
Cellular process
Regulation of cellular component organization
Cellular component organization
Intracellular signal transduction
Cell adhesion
Groups at similarity 0.5
Signal > 0.5
FDR
1.0e-14
1.0e-13
1.0e-11
1.0e-10
1.0e-08
1.0e-07
Gene count
500
2000
3000
4500

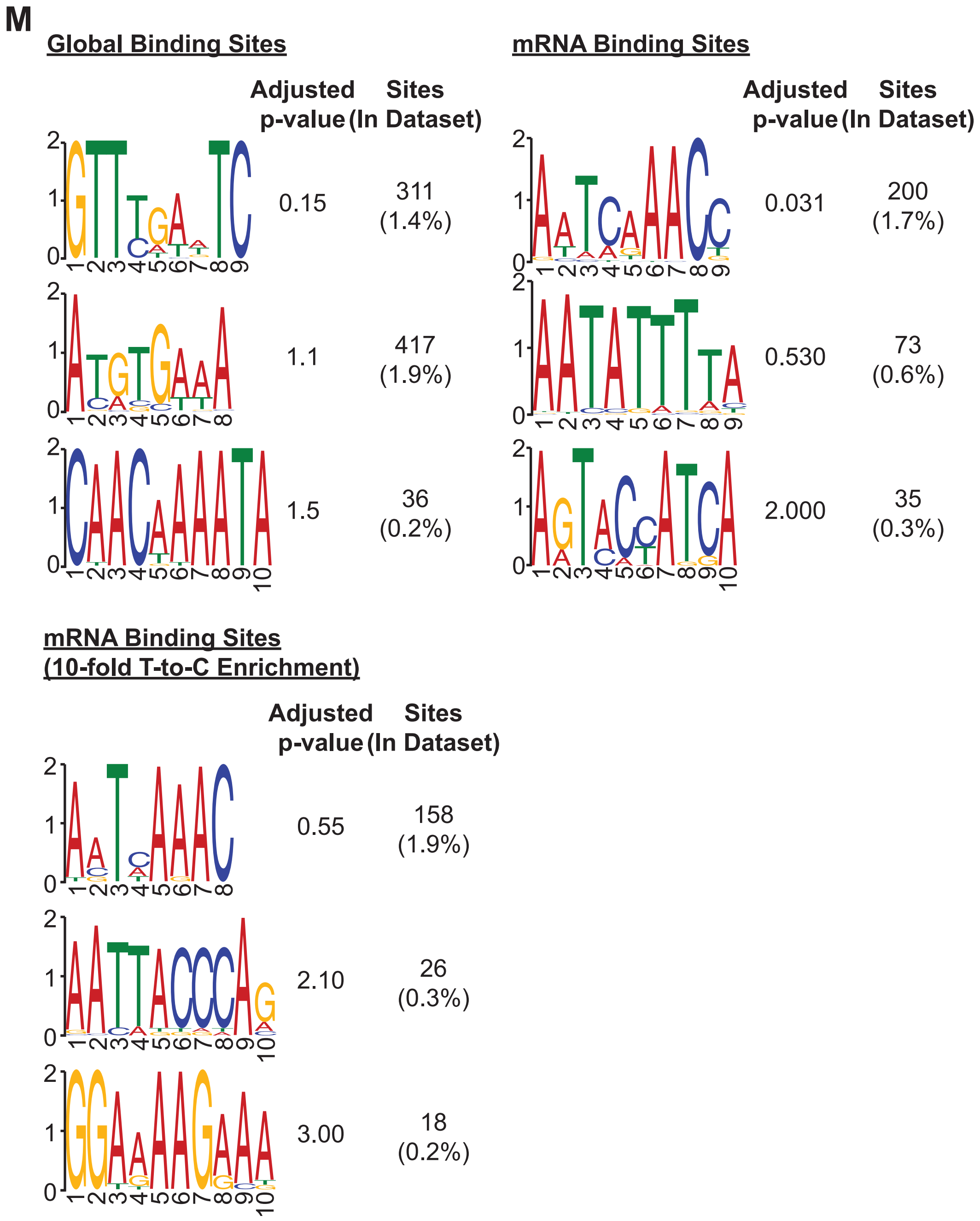
M
Global Binding Sites
Adjusted p-value
Sites (In Dataset)
0.15
311 (1.4%)
1.1
417 (1.9%)
1.5
36 (0.2%)
mRNA Binding Sites
Adjusted p-value
Sites (In Dataset)
0.031
200 (1.7%)
0.530
73 (0.6%)
2.000
35 (0.3%)
mRNA Binding Sites (10-fold T-to-C Enrichment)
Adjusted p-value
Sites (In Dataset)
0.55
158 (1.9%)
2.10
26 (0.3%)
3.00
18 (0.2%)

### Supplemental Figure 3.

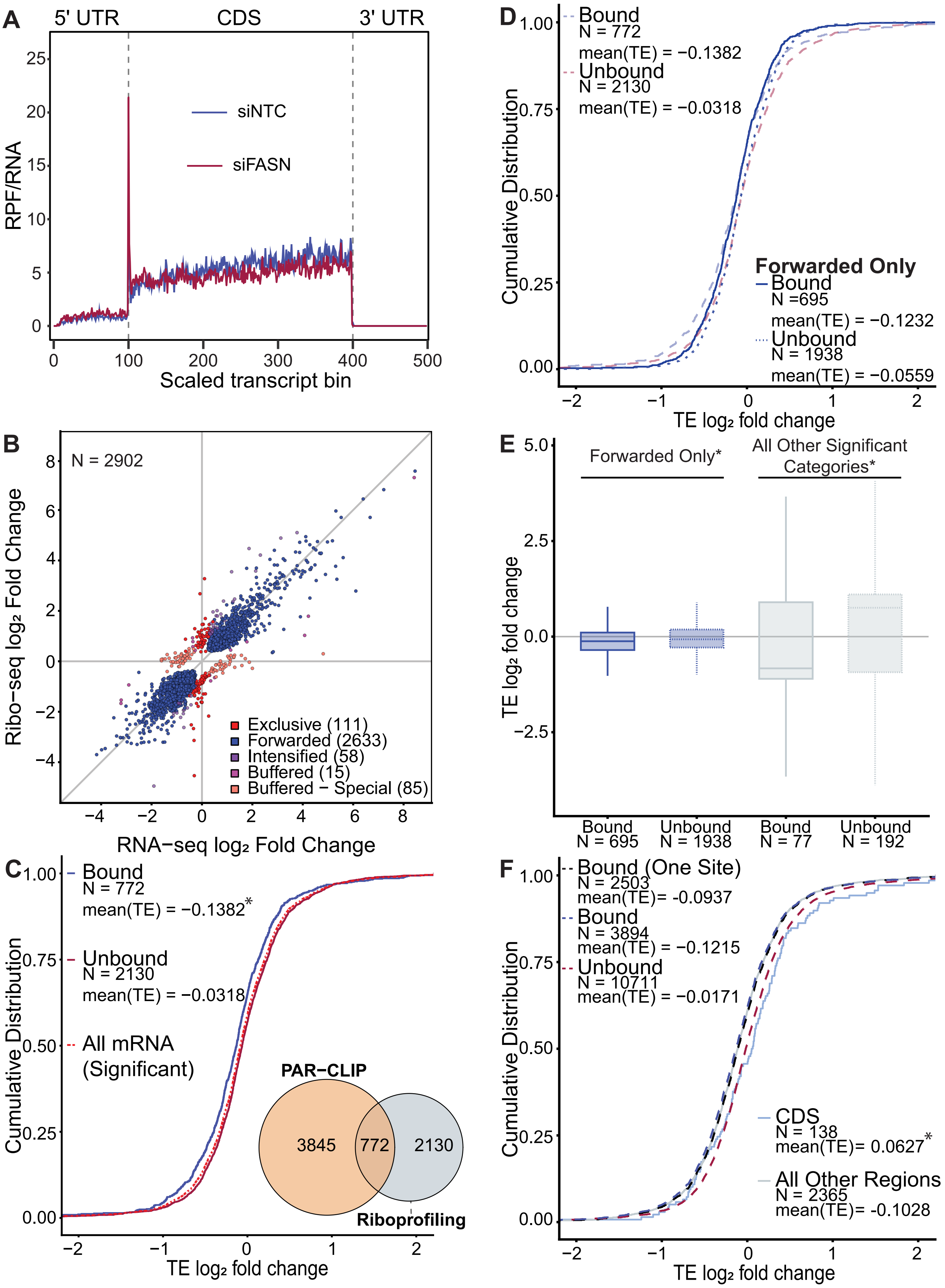

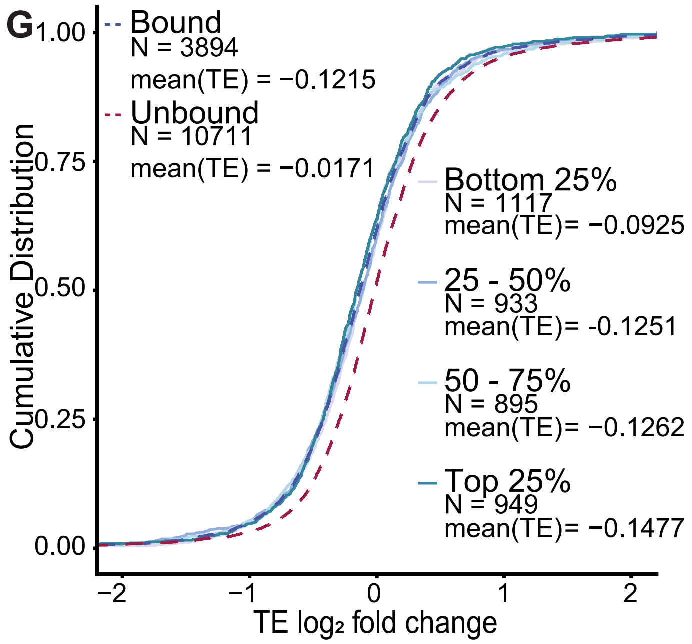
G
Bound
N = 3894
mean(TE) = −0.1215
Unbound
N = 10711
mean(TE) = −0.0171
Bottom 25%
N = 1117
mean(TE)= −0.0925
25 - 50%
N = 933
mean(TE)= -0.1251
50 - 75%
N = 895
mean(TE)= −0.1262
Top 25%
N = 949
mean(TE)= −0.1477
Cumulative Distribution
1.00
0.75
0.50
0.25
0.00
−2
−1
0
1
2
TE log2 fold change

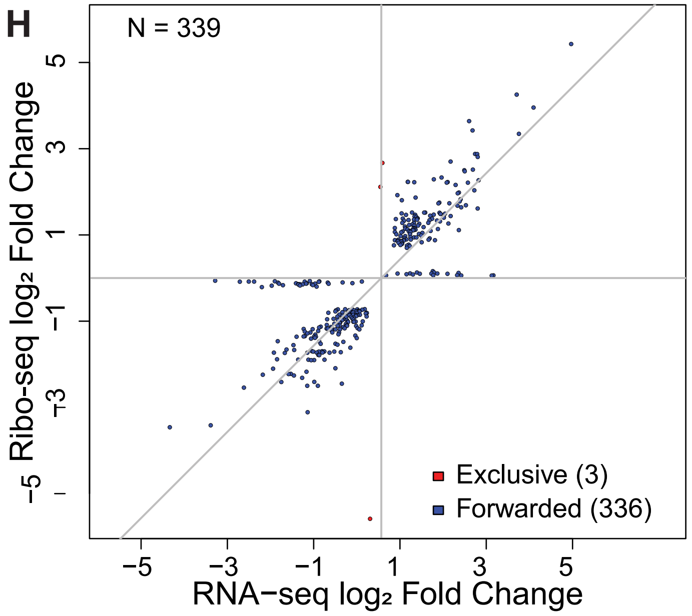
H
N = 339
Exclusive (3)
Forwarded (336)
Ribo-seq log2 Fold Change
5
3
1
−1
−3
−5
−5
−3
−1
1
3
5
RNA−seq log2 Fold Change

#### Supplemental Figure 4.

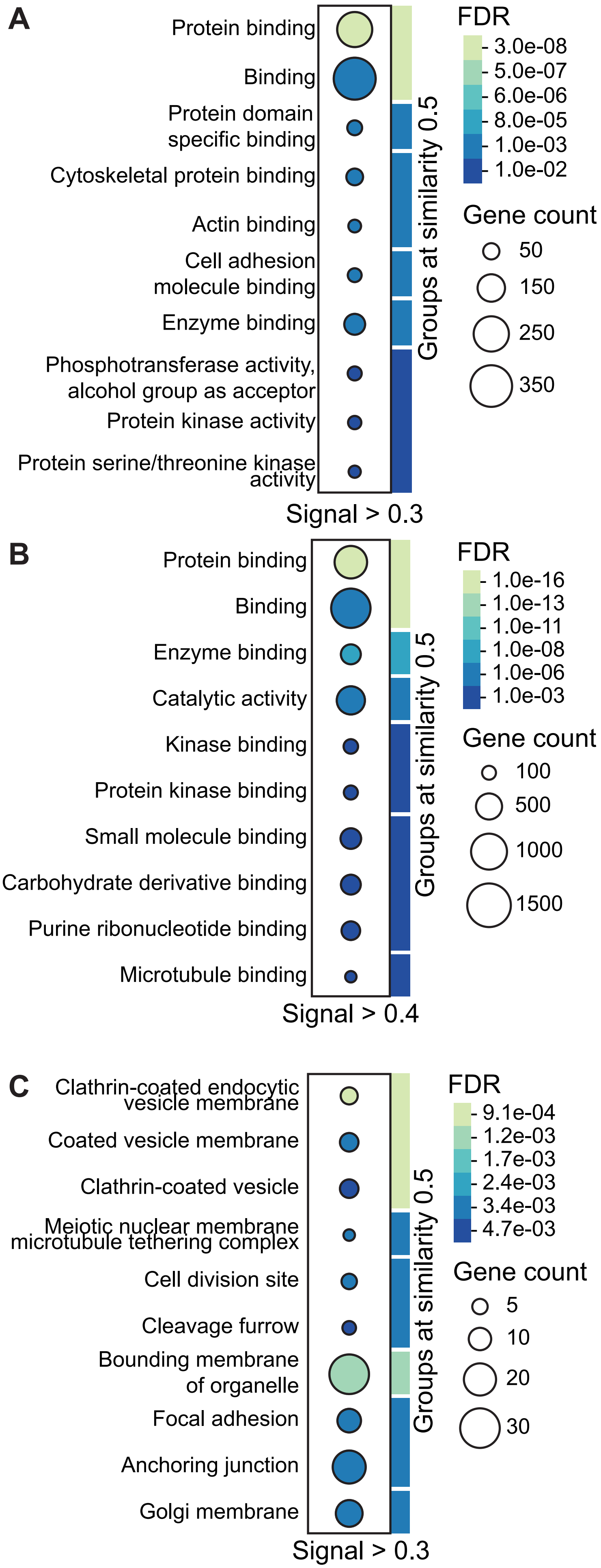
